# Assessing the Involvement of Platelet Degranulation in the Therapeutic Properties of Exosome Derived from Amniotic Epithelial Cells through Enrichment and Interaction Network Analysis

**DOI:** 10.1101/2021.01.07.425685

**Authors:** M. Valizadeh, A. Haider Bangash, D. Hayati, A. Jafari, H. Rajabi-Maham

## Abstract

Platelet degranulation allows the release of large secretable pools of biologically active proteins which are critical in wound healing initiation and angiogenesis. Exosomes, which can transport a diverse suite of macromolecules, derived from amniotic epithelial cells (AEC-Exo) improve wound healing and angiogenesis. However, the underlying mechanisms are still unclear. In this investigation, we performed a user-friendly bioinformatics analysis system to identify association among the angiogenic and wound healing effects of AEC-Exo treatments. To this end, FunRich software was used, and linked to the Universal Protein Resource (UniProt) as a background database. Several enrichment analyses, including biological process, cellular component, molecular function, and protein domains were conducted on AEC-Exo proteome. Furthermore, to identify the proteins involved in platelet degranulation and evaluate protein–protein association information, comparative analyses and interaction network analyses were illustrated using the NCBI BioSystems, ExoCarta, and STRING databases. Our results indicated the statistically significant association between the proteome in AEC-Exo, platelet degranulation, and their corresponding processes. Therefore, the involvement of platelet degranulation in AEC-Exo proteins may elucidate the angiogenic and wound-healing effects of AEC-Exo treatments.

## 1 Introduction

The wound-healing process is a complex mechanism that requires angiogenesis as a process in which new blood vessels are formed from pre-existing ones (1). At the site of injury, platelets are essential in regulating angiogenesis but their role in repairing damaged vessels during the wound healing process is still not well known (2). Platelet degranulation allows the release of large secretable substances that promote tissue repair, angiogenesis, and inflammation including Vascular Endothelial Growth Factors (VEGFs), Basic Fibroblast Growth Factors (FGF-2), and Platelet Derived Growth Factors (PDGFs), Insulin-Like GF (IGF1), Transforming GF-beta 1 (TGF-β1), Vascular Endothelial GF (VEGF), Basic Fibroblastic GF (bFGF), and Epidermal GF (EGF) (3).

Growing evidence has suggested that the therapeutic efficacy of placental cells might depend on paracrine mechanisms, through the secretion of Extracellular vesicles (4). In this regard, small lipid-bound(30-150 nm), cellularly secreted vesicles that are called exosomes have especially attracted the attention of researchers (5). Exosomes are multivesicular bodies (MVBs) matured from early endosomes. They mediate local and systemic intercellular communication by fusing with target cells and releasing their macromolecular contents. Exosomes play an important role in many biological processes, including immune response and inflammation, blood coagulation, and angiogenesis (6, 7). Several studies have shown that exosomes derived from amniotic epithelial cells (AEC-Exo) improve wound healing and angiogenesis but their primary mechanisms remain unclear (8, 9).

In order to get a comprehensive understanding of the association between the angiogenic and wound healing effects of AEC-Exo treatments in current studies, we utilized a user-friendly bioinformatics analysis system. FunRich is a software with added options for functional enrichment and interaction network analyses of the genes and proteins (10). The Universal Protein Resource (UniProt) was examined via the Funrich tool to conduct enrichment analyses, including Gene ontology and protein domains. Moreover, in order to identify the proteins involved in degranulation and assess protein-protein association information, comparative and interaction network analyses were performed using various databases.

## 2 Method

### 2.1 AEC-Exo proteins data

The AEC-Exo proteins data of a proteomics analysis, deposited by Sheller et al. (2016), was downloaded from https://journals.plos.org/ (11). For further analyses, all protein ID reported in the study were converted to the UniProt protein identifier, and the repeated proteins were removed.

### 2.2 Enrichment analyses

The functional enrichment analyses of AEC-Exo proteins were performed using FunRich software version 3.1.3 available for public access. FunRich users are provided a user-friendly GUI (graphical user interface) allowing them to download directly from the FunRich database and UniProt, in order to further analyze their data. Therefore, a database from UniProt for human proteins (Taxon ID: 9606, ftp://ftp.uniprot.org/) was downloaded and the AEC-Exo proteins were analyzed against it. Several options were used for the enrichment analysis, including biological process, cellular component, molecular function, and protein domains. Subsequently, the results of the analyses were depicted graphically in the form of a column chart.

### 2.3 Comparative analyses

In order to find the proteins involved in degranulation, Funrich was used to generate a Scalable-Venn diagram and Matrix table (also known as pair-wise comparison) among the total platelet degranulation proteins, the AEC-Exo proteins, and the human exosomal proteins. One hundred and twenty-five platelet degranulation proteins were chosen in the NCBI BioSystems database via the Swiss-Prot/Uniprot filter, and all selected protein data was downloaded from the NCBI protein database. The human exosomal protein data was subsequently downloaded from ExoCarta (http://exocarta.org/download). Finally, to quantify the relative expression of proteins identified in the Human Proteome Map, the heatmap image was generated

### 2.4 Interaction network analysis

The STRING database (http://string-db.org/) was applied to illustrate and analyze protein-protein interaction (PPI) network. The interaction network analysis of FunRich was constructed for AEC-Exo proteins using UniProt as a background database.

## 3 Result

### 3.1 AEC-Exo proteins and platelet degranulation

Biological processes and Reactome pathway analysis indicated that 20.2% and 16.6% of the proteins in AEC-Exo are remarkably engaged in platelet degranulation. (p<0.001, fold change = 17, 22.1) (Figure 1, 2). In addition, 13.5% of these proteins are engaged in the regulation of Insulin-like growth factor (IGF) transport and its uptake by insulin-like growth factor-binding proteins (IGFBPs) (fold change = 11.7).

**Figure 1:**
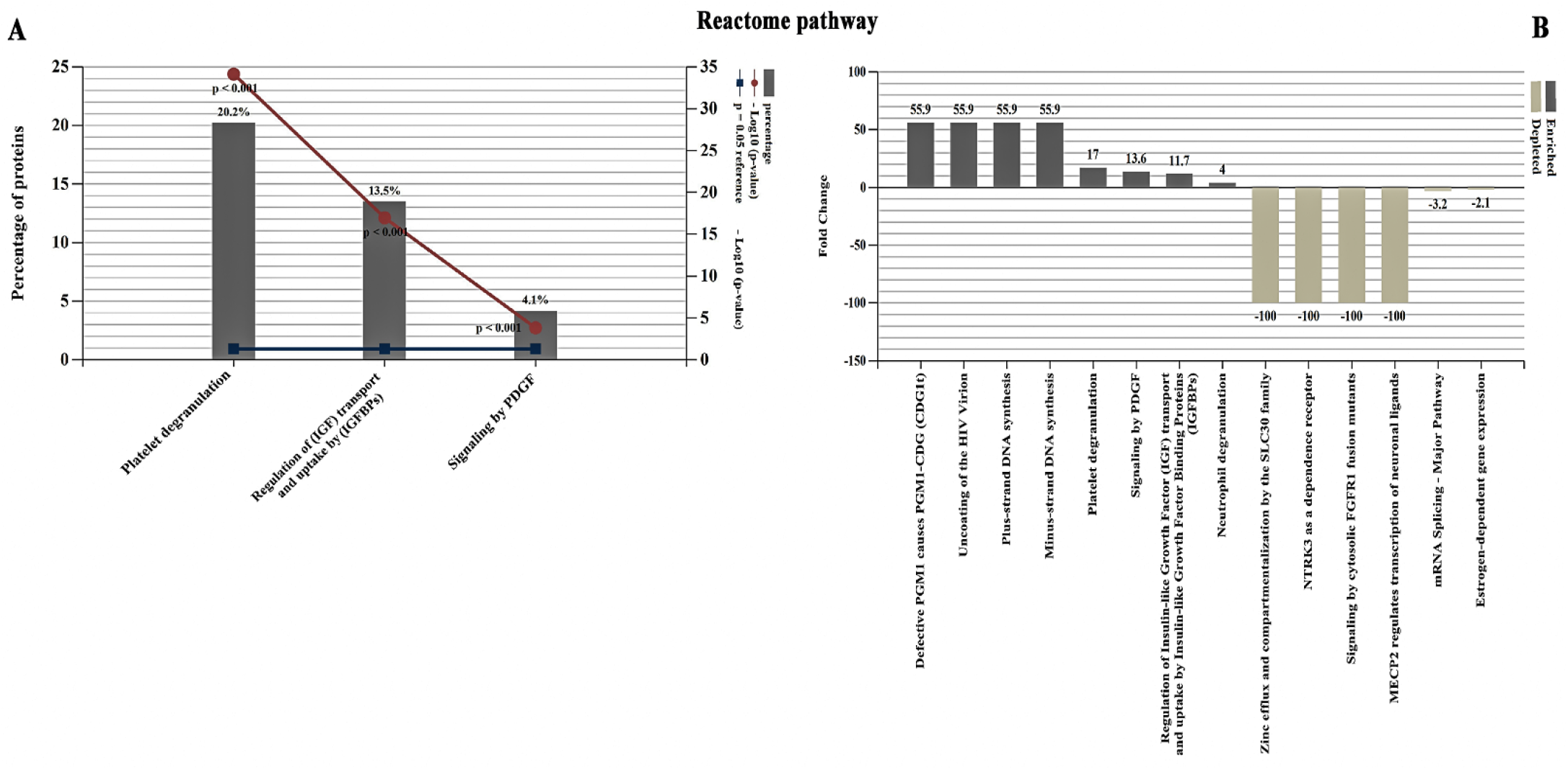
Reactome pathway analysis; 20.2 % of the proteins in AEC-Exo are engaged in platelet degranulation (p<0.001, fold change = 17)

**Figure 2:**
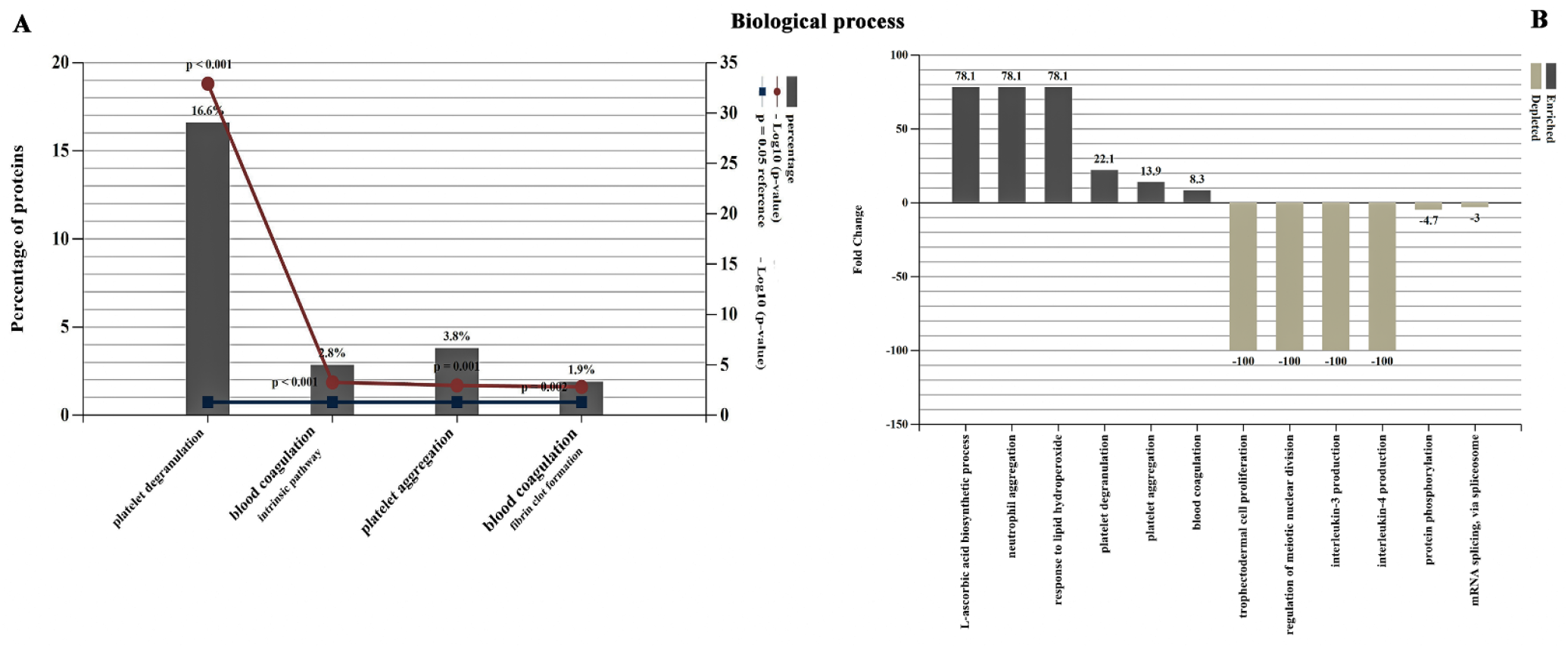
Biological process analysis; 16.6% of the proteins in AEC-Exo are involved in platelet degranulation (p<0.001, fold change = 22.1)

### 3.2 Protein domain, molecular function and cellular component analysis

Protein domain analysis revealed the percentage of the proteins in AEC-Exo, including epidermal growth factor (EGF), calcium-binding EGF (EGF-CA), trypsin-like serine protease (Tryp-SPc), and serine proteinase inhibitors (SERPIN) (Fig. 3). The molecular function analysis of the proteins further revealed the involvement of calcium ion binding, platelet-derived growth factor binding and signaling receptor binding proteins (Fig. 4). The cellular component analysis of the proteins in AEC-Exo also showed the percentage of blood microparticle, platelet alpha granule lumen, platelet dense granule lumen and platelet alpha granule proteins (p<0.001) (Fig. 5).

**Figure 3:**
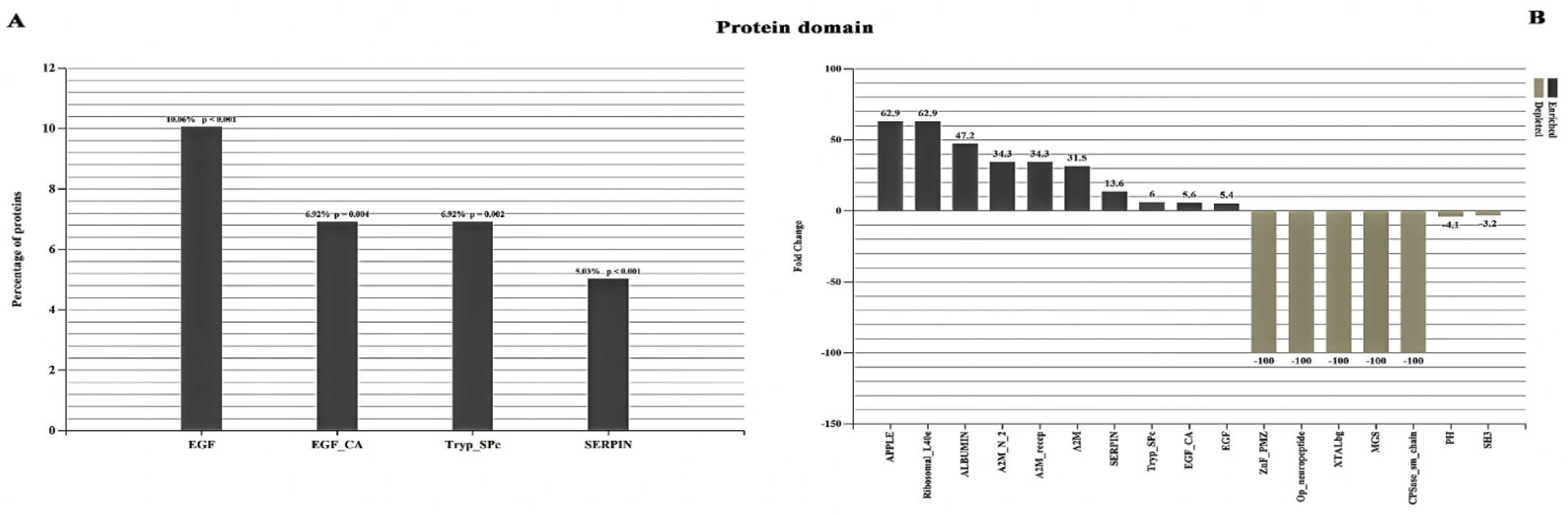
(A) Protein domain analysis of the proteins in AEC-Exo (p<0.001) (B) Separate number of fold enriched/depleted of every protein domain

**Figure 4:**
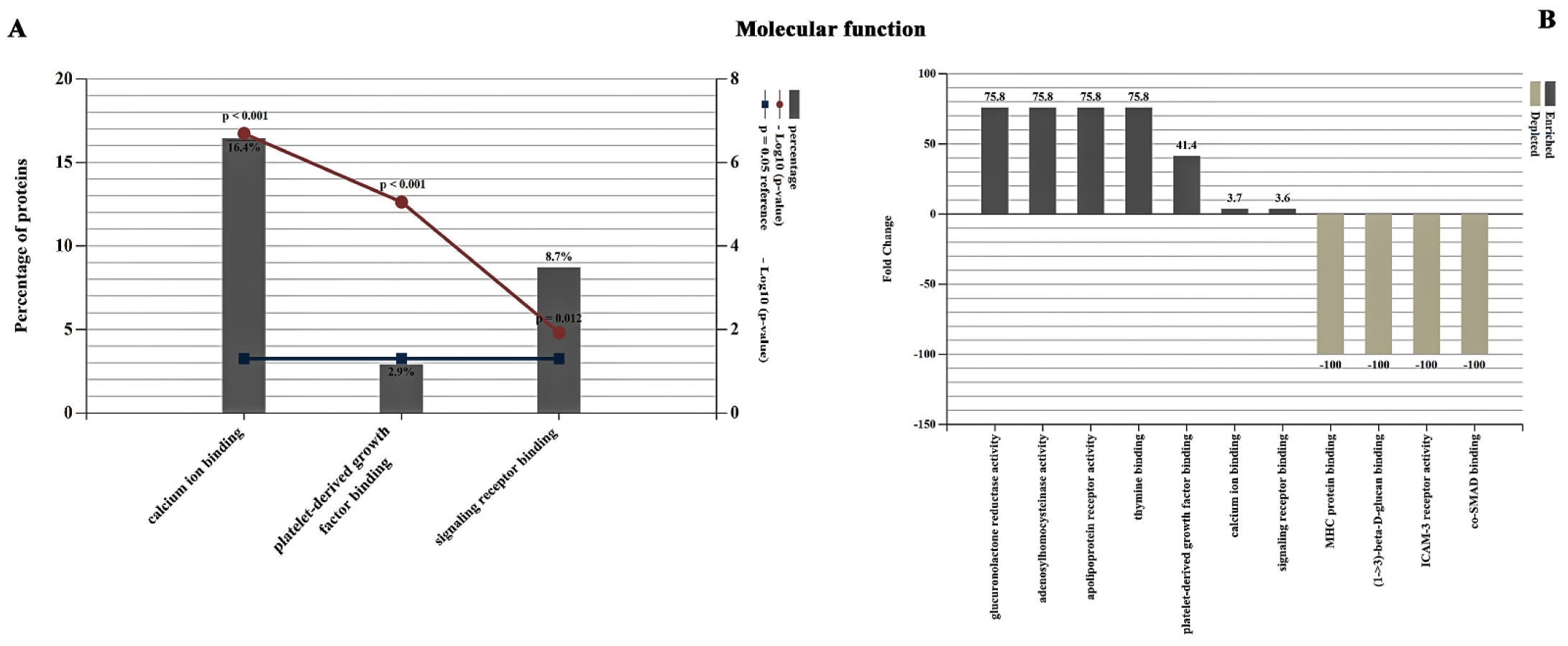
(A) Molecular function analysis of the AEC-Exo proteins (B) Separate number of fold enriched/depleted of each molecular function

**Figure 5:**
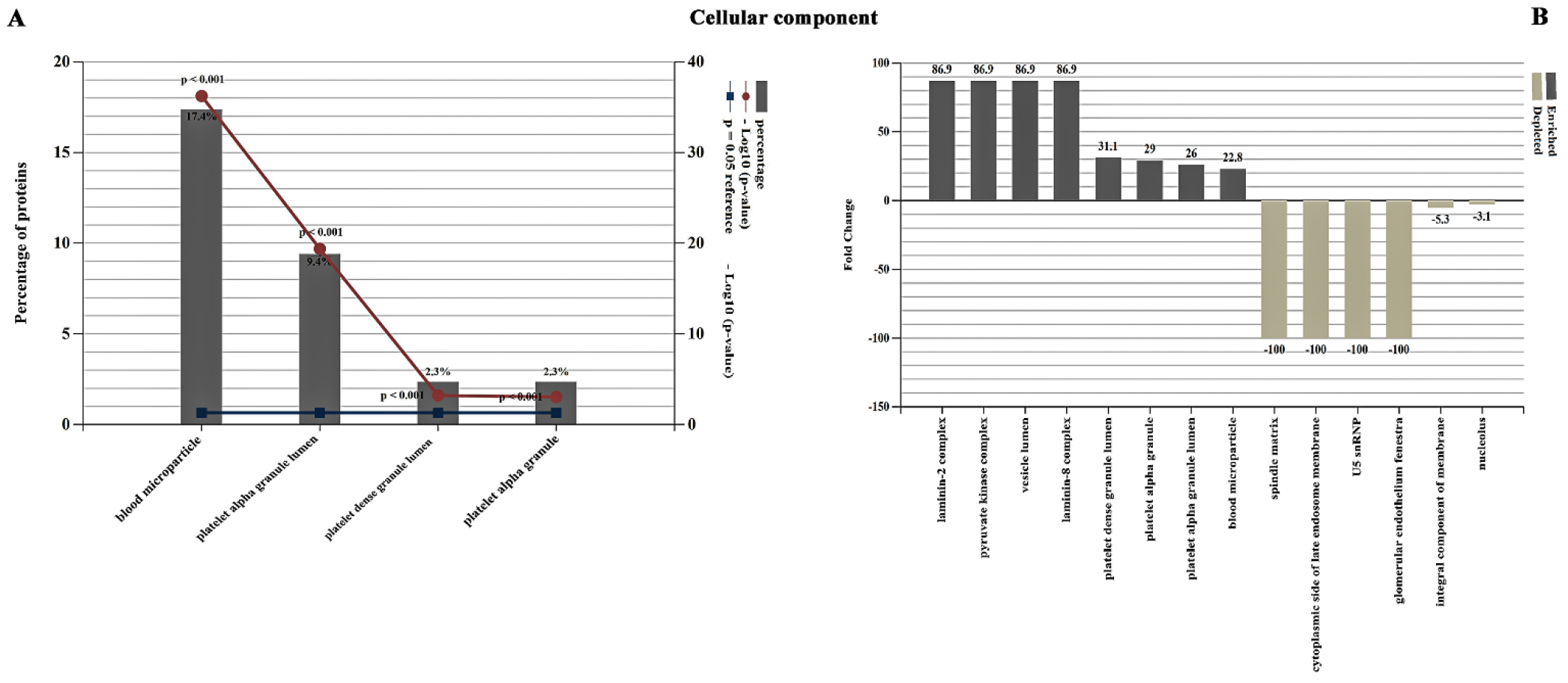
(A) Cellular component analysis of the AEC-Exo proteins (B) Separate number of fold enriched/depleted of cellular component

### 3.3 Protein identification and comparison analysis

Thirty-seven proteins were identified that were involved in platelet degranulation in the AEC-Exo (Fig. 6, Table 1). The Venn diagram and the Matrix table revealed the overlap of 103 proteins between human exosomal proteins, and total platelet degranulation proteins. Figure 7 illustrated the heatmap image of AEC-Exo proteins involved in platelet degranulation based on the human proteome map.

**Figure 6:**
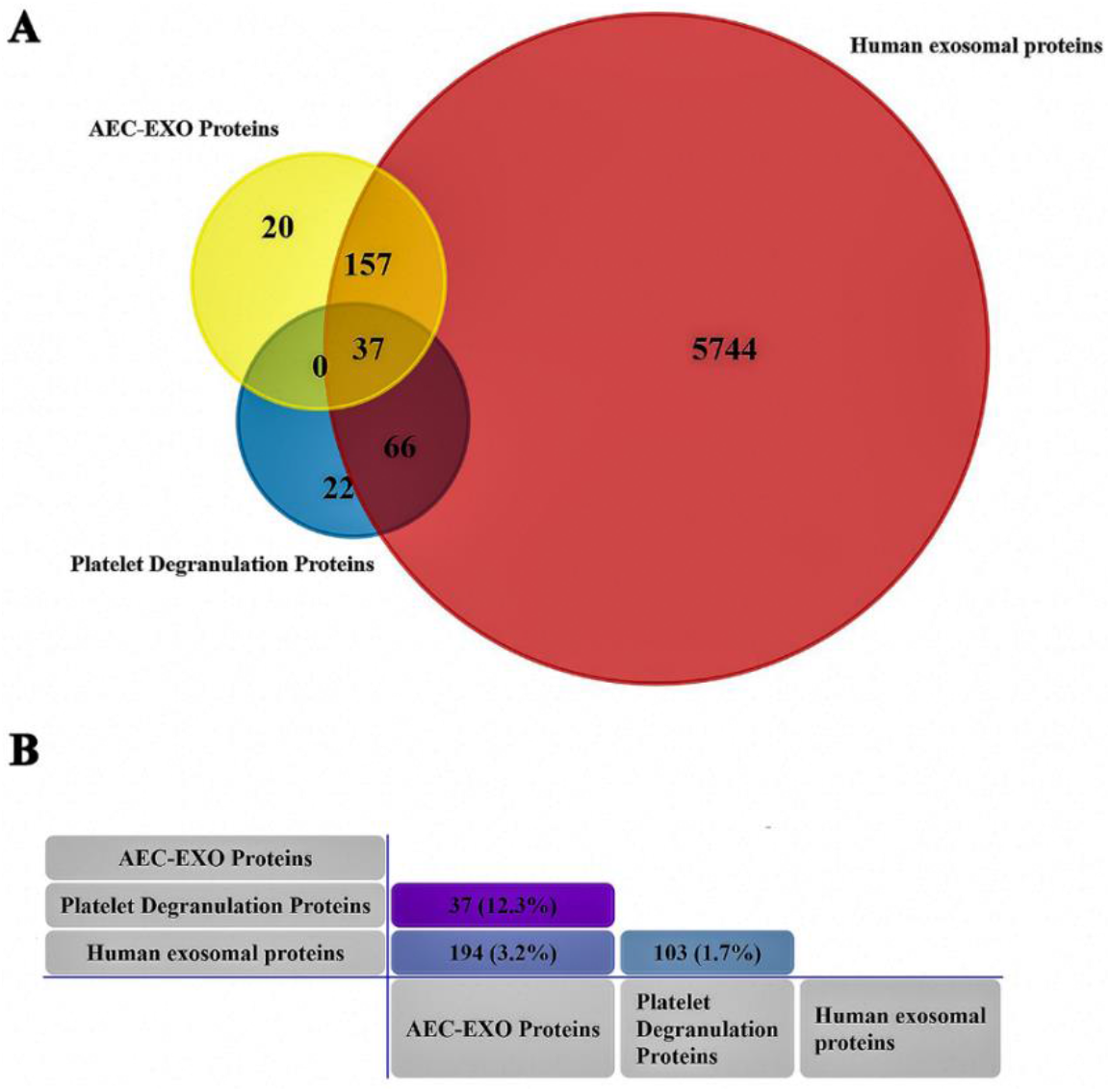
(A) Venn diagram showing the overlaps between total platelet degranulation proteins, human exosomal proteins and AEC-Exo. (B) Matrix table (pair-wise comparison) for AEC-Exo, total platelet degranulation proteins and human exosomal proteins.

**Table 1:** The List of the proteins in AEC-Exo involved in platelet degranulation

| Identifier | Protein names | Length (aa) |
| --- | --- | --- |
| A2M | Alpha-2-macroglobulin | 1474 |
| ACTN1 | Alpha-actinin-1 | 914 |
| ACTN4 | Alpha-actinin-4 | 911 |
| ALDOA | Fructose-bisphosphate aldolase A | 418 |
| APOA1 | Apolipoprotein A-I | 267 |
| APOH | Beta-2-glycoprotein 1 | 345 |
| CAP1 | Adenylyl cyclase-associated protein 1 | 475 |
| CD109 | CD109 antigen | 1445 |
| CLEC3B | Tetranectin | 202 |
| CLU | Clusterin | 449 |
| Cofilin-1 | Cofilin-1 | 166 |
| F13A1 | Coagulation factor XIII A chain | 732 |
| F5 | Coagulation factor V | 2224 |
| FERMT3 | Fermitin family homolog 3 | 667 |
| FGB | Fibrinogen beta chain | 491 |
| FGG | Fibrinogen gamma chain | 453 |
| FLNA | Filamin-A | 2647 |
| IGF2 | Insulin-like growth factor II | 236 |
| ITIH3 | Inter-alpha-trypsin inhibitor heavy chain H3 | 890 |
| ITIH4 | Inter-alpha-trypsin inhibitor heavy chain H4 | 930 |
| PFN1 | Profilin-1 | 140 |
| PLG | Plasminogen | 810 |
| PPIA | Peptidyl-prolyl cis-trans isomerase A | 165 |
| PROS1 | Vitamin K-dependent protein S | 676 |
| QSOX1 | Sulfhydryl oxidase 1 | 747 |
| SERPINE1 | Plasminogen activator inhibitor 1 | 402 |
| SERPINF2 | Alpha-2-antiplasmin | 491 |
| TAGLN2 | Transgelin-2 | 220 |
| TF | Serotransferrin | 698 |
| THBS1 | Thrombospondin-1 | 1170 |
| TIMP3 | Metalloproteinase inhibitor 3 | 211 |
| TLN1 | Talin-1 | 2541 |
| TTN | Titin | 35991 |
| TUBA4A | Tubulin alpha-4A chain | 448 |
| VCL | Vinculin | 1134 |
| VWF | Von Willebrand factor | 2813 |
| WDR1 | WD repeat-containing protein 1 | 606 |

**Figure 7:**
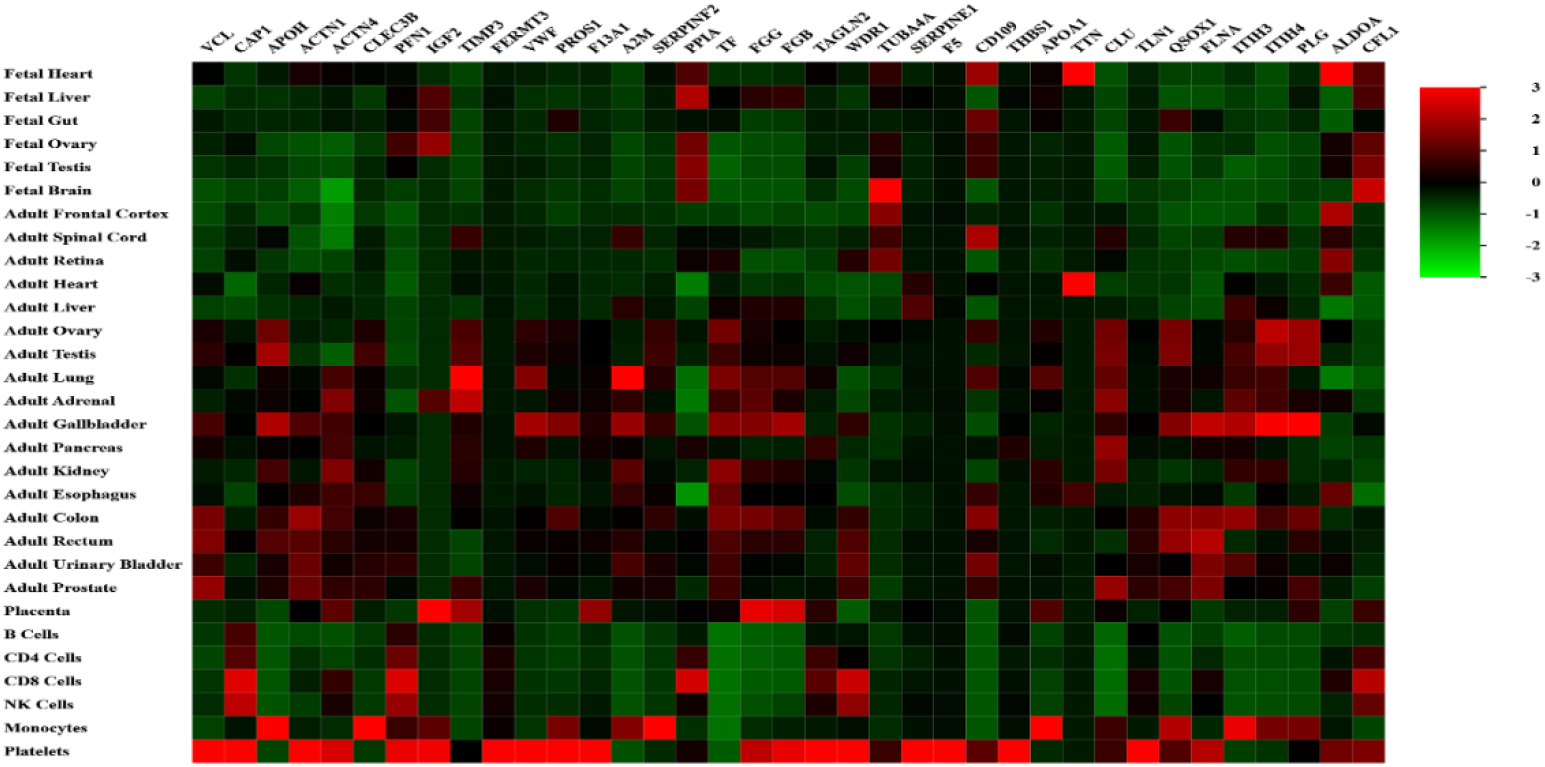
Heat map representing of the proteins in AEC-Exo involved in platelet degranulation (green–lowest abundance and red–highest abundance)

### 3.4 Predicting PPI network of platelet degranulation

To investigate the interactions between the proteins that participated in platelet degranulation, a PPI network was constructed. The interaction network planetary layout of AEC-EXO proteome data via the FunRich displayed 39 genes significantly enriched for platelet degranulation (p<0.001) (Fig. 8). Fibronectin (FN1) with the highest degree of distribution is placed in the center of the network construction. On the other hand, the number of 259 edges through the STRING represents protein-protein associations that are significant, therefore the proteins contribute to a shared function equally (PPI enrichment p-value < 1.0e-16) (Fig. 9).

**Figure 8:**
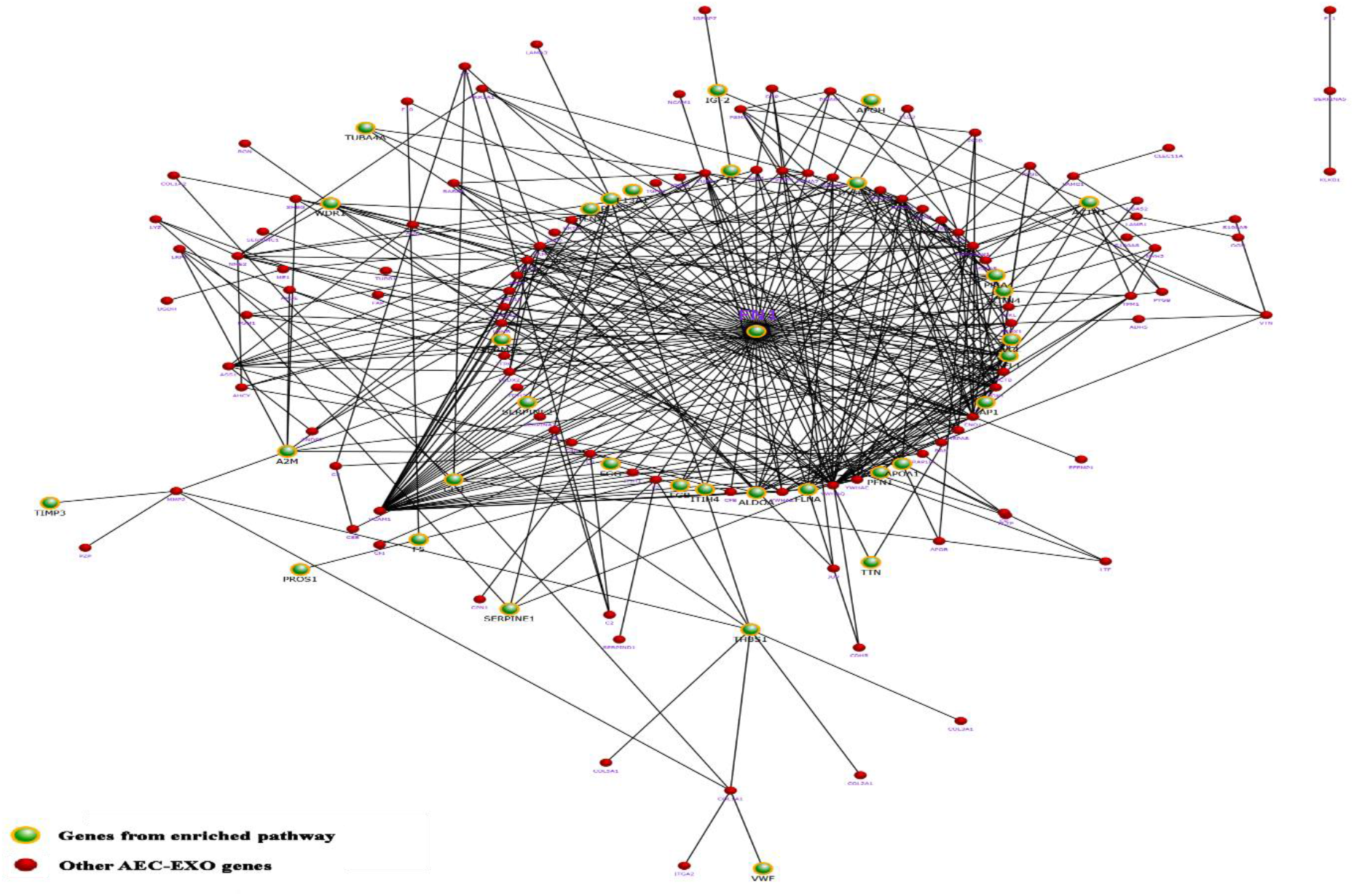
Interaction network planetary layout of AEC-Exo proteome data via the FunRich (p<0.001). The list of genes enriched in platelet degranulation were highlighted within the interactions network.

**Figure 9:**
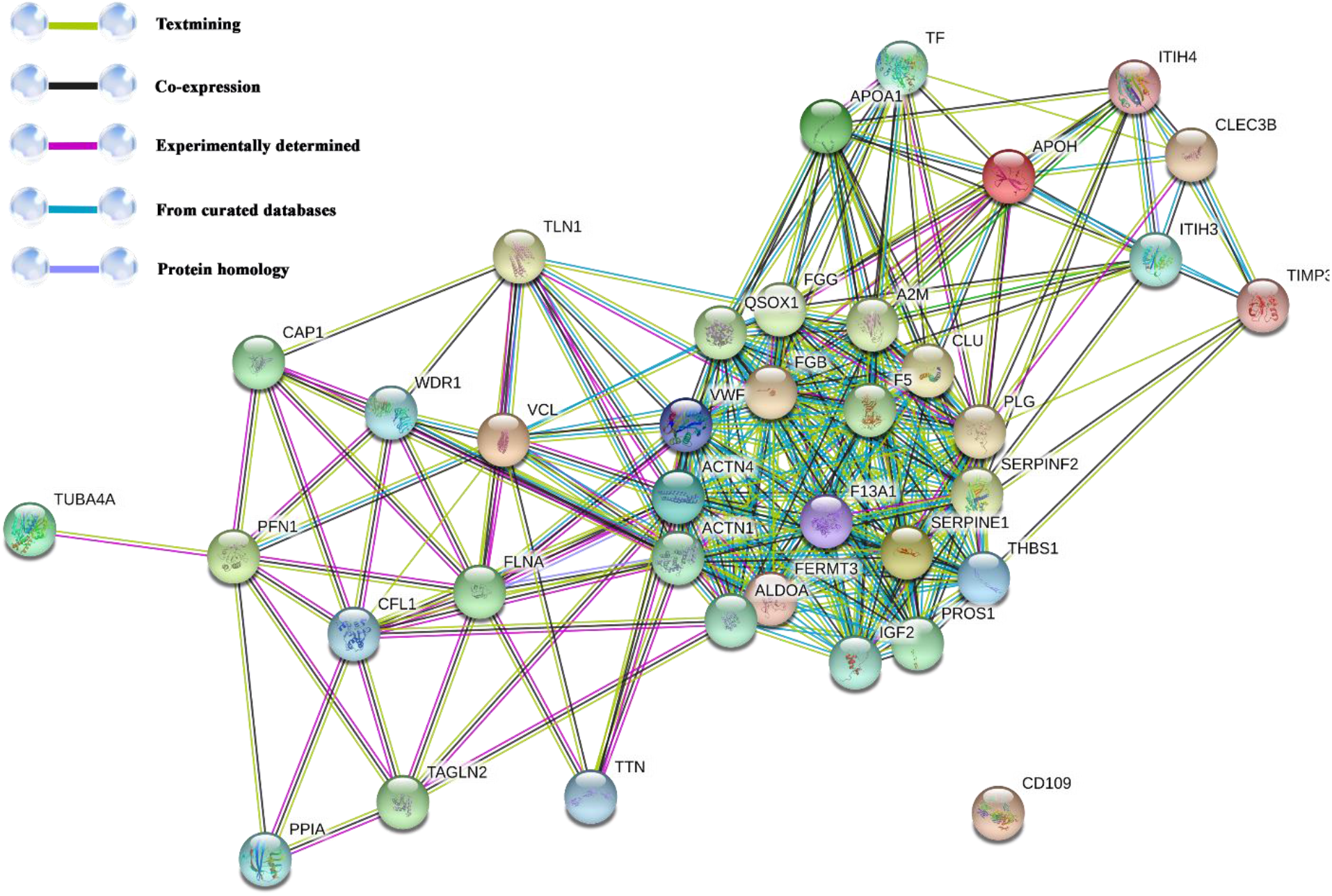
PPI network analysis with the STRING database of the 37 proteins in AEC-Exo involved in platelet degranulation. The edges represent protein-protein associations. (PPI enrichment p-value < 1.0e-16)

## 4 Discussion

The results of the present investigation demonstrate the significant association between AEC-Exo proteome, platelet degranulation, and their corresponding processes. A significant relevance was observed between the proteins in AEC-Exo, and platelet degranulation which is pivotal as a primary mechanism for the healing process and angiogenesis. In addition, interaction network analysis using the STRING and FunRich validated enrichment in the platelet degranulation. Over the past decades, efforts have been made to modulate angiogenesis as a therapeutic strategy to either increase revascularization of ischemic tissues or inhibit angiogenesis in cancer, ocular, joint, or skin disorders. Consistent with the literature, other studies have confirmed AEC-Exo and placental exosome’s role in increasing wound healing and angiogenesis (12–14). Consequently, the involvement of platelet degranulation in AEC-Exo proteins may illustrate the angiogenic effects of AEC-Exo treatments in current studies.

The results of protein domain analysis showed that the highest protein percentage belongs to EGF domain. Each EGF domain is comprised about 40 to 45 amino acids and found in a large variety of proteins. Most of the angiogenic factors contain the EGF-like domain, which may exhibit a mitogenic effect on endothelial cells and facilitate neovascularization (15, 16). For instance, Kao YC et al. (2012) demonstrated that the EGF-like domain is essential for CD93 to induce angiogenesis (17). Hence, the inclusion of these domains in the AEC-EXO proteins increasingly affects the modulation of angiogenesis and wound healing. The Reactome pathway analysis revealed the importance of the IGFBP family in the AEC-EXO proteins. It has been shown that they have both stimulate and inhibit angiogenesis (18). Activation of VEGF is modulated by IGFBP-2 to induce an angiogenic-type response in human umbilical vein endothelial cells (HUVEC) in vitro. Studies have shown IGF1 stimulates endothelial cell sprouting in an in vitro assay. PDGF has an important role in angiogenesis but molecular component analysis of the proteins in AEC-Exo only showed four percent participation (19–21).

Based on molecular function analysis conducted in this study, the proteins in AEC-Exo remarkably participated in calcium ion binding, platelet-derived growth factor binding, and signaling receptor binding. Bioactive molecules of dense granules containing catecholamines, histamine, serotonin, ADP, ATP, calcium ions, and dopamine, which are active in vasoconstriction, increased capillary permeability, attracting and activating macrophages, tissue modulation, and regeneration. The secretion of PDGFB, TGF-beta1, bFGF, and VEGF is considerably mediated by the amount of calcium and thrombin (Thr) added to the platelet-rich plasma (PRP), and PRP supernatants are more mitogenic for endothelial cells than whole-blood supernatants. Endothelial cells are crucial in vessel formation by angiogenesis and vasculogenesis. Intracellular calcium signals are involved in different critical phases in the regulation of the complex and multi-stepped process of angiogenesis. Furthermore, regarding either external activation with calcium and thrombin or internal activation with endogenous tissue thromboplastin, the fibrin matrix forms by polymerization of plasma fibrinogen (22–24). Accordingly, our observations further support and strengthen the importance of the AEC-Exo in angiogenesis and wound healing by platelet degranulation.

## 5 Conclusion

Based on these findings, the involvement of platelet degranulation in AEC-Exo proteins may elucidate the angiogenic and wound healing effects of AEC-Exo treatments in current studies. Collectively, our bioinformatic-based study provided the AEC-EXO proteome, which can be of high interest to reveal biological insights about the healing process and angiogenesis.

